# Brevibacillin 2V, a novel antimicrobial lipopeptide with an exceptional low hemolytic activity

**DOI:** 10.1101/2021.04.11.439345

**Authors:** Xinghong Zhao, Xiaoqi Wang, Rhythm Shukla, Raj Kumar, Markus Weingarth, Eefjan Breukink, Oscar P. Kuipers

**Affiliations:** Department of Molecular Genetics, Groningen Biomolecular Sciences and Biotechnology Institute, University of Groningen, Nijenborgh7, 9747 AG Groningen, The Netherlands; Membrane Biochemistry and Biophysics, Bijvoet Centre for Biomolecular Research, Department of Chemistry, Faculty of Science, Utrecht University, Padualaan 8, Utrecht, The Netherlands; NMR Spectroscopy, Bijvoet Centre for Biomolecular Research, Department of Chemistry, Faculty of Science, Utrecht University, Padualaan 8, Utrecht, The Netherlands

**Keywords:** Antimicrobial activity, Lipopeptide, NRPs, Brevibacillin, *Brevibacillus*, *Enterococcus*, *Staphylococcus*

## Abstract

Bacterial non-ribosomally produced peptides (NRPs) form a rich source of antibiotics, including more than twenty of these antibiotics that are used in the clinic, such as penicillin G, colistin, vancomycin, and chloramphenicol. Here we report the identification, purification, and characterization of a novel NRP, i.e., brevibacillin 2V, from *Brevibacillus laterosporus* DSM 25. Brevibacillin 2V has strong antimicrobial activity against Gram-positive bacterial pathogens, including difficult-to-treat antibiotic-resistant *Enterococcus faecium, Enterococcus faecalis*, and *Staphylococcus aureus*. Notably, brevibacillin 2V has a much lower hemolytic activity to eukaryotic cells than previously reported NRPs of the lipo-tridecapeptide family, including other brevibacillins, which makes it a promising candidate for antibiotic development. In addition, our results demonstrate that brevibacillins display synergistic action with established antibiotics against Gram-negative bacterial pathogens. Thus, we identified and characterized a novel and promising antibiotic candidate (brevibacillin 2V) with a low hemolytic activity, which can be used either on its own or as a template for further total synthesis and modification.

## Introduction

The non-ribosomally produced peptides (NRPs) have formed a rich source of antimicrobials, including more than twenty marketed antibiotics, such as chloramphenicol, penicillin G, colistin, and vancomycin (Süssmuth and Mainz, 2017). Among these, lipopeptides form a rich class of NRPs that have shown strong antimicrobial activity against Gram-positive (Hamamoto et al., 2015; Ling et al., 2015) and/or Gram-negative (Cochrane et al., 2014; Cociancich et al., 2015) pathogens.

Large numbers of antimicrobial NRPs have been isolated from different *Brevibacillus* spp (Yang and Yousef, 2018), including many linear lipopeptides, such as bogorol A-E (Barsby et al., 2001, 2006), brevibacillin (Yang et al., 2016), and brevibacillin V (Wu et al., 2019). Bogorol A-E, brevibacillin, and brevibacillin V are produced by *Brevibacillus laterosporus* PNG-276, *Brevibacillus laterosporus* OSY-I1, and *Brevibacillus laterosporus* fmb70 (Barsby et al., 2001, 2006; Yang et al., 2016; Wu et al., 2019), respectively, and all of them can be grouped into the family of non-ribosomally produced linear lipo-tridecapeptides (Yang and Yousef, 2018). These lipo-tridecapeptides show strong antimicrobial activity against Gram-positive pathogenic bacteria, including methicillin-resistant *Staphylococcus aureus* and vancomycin-resistant *Enterococcus spp*. (Barsby et al., 2006; Yang et al., 2016; Wu et al., 2019). However, many lipo-tridecapeptides have the disadvantage of being highly hemolytic (Li et al., 2018).

In this study, two novel lipo-tridecapeptides, brevibacillin I and brevibacillin 2V, were discovered from *Brevibacillus laterosporus* DSM 25 by genome mining, and two of the known lipo-tridecapeptides, brevibacillin and brevibacillin V, were also isolated from the same strain. All these four lipo-tridecapeptides showed strong antimicrobial activity against Gram-positive pathogenic bacteria, and all of these four lipo-tridecapeptides showed synergistic effects with marketed antibiotics against Gram-negative pathogens. Notably, only brevibacillin 2V did not exhibit hemolytic activity when present at 128 µg/mL, demonstrating its potency to be further studied and developed as a candidate antibiotic for controlling specific antibiotic-resistant bacterial pathogens.

## Materials and Methods

### Bacterial Strains Used and Growth Conditions

Strains used in this study are listed in Supplementary Table 1. *Brevibacillus laterosporus* DSM 25 cells were inoculated on in LB and incubated at 37 °C for prepare overnight culture. For production of brevibacillins, an overnight culture of *Brevibacillus laterosporus* DSM 25 cells was inoculated (50-fold dilution) in minimal expression medium (MEM) and grown 36 h at 30 ^0^C. All indicator strains were inoculated on LB and incubated at 37 °C for preparing the overnight cultures.

### Purification of Cationic Peptides from a *Brevibacillus laterosporus* DSM25 Culture

An overnight culture of *Brevibacillus laterosporus* DSM 25 cells was inoculated in MEM and grown 36 h at 30 ^0^C. Subsequently, the culture was centrifuged at 15,000g for 15 min, and the supernatant was collected and adjusted the pH to 7.2. After that, the culture was applied to a CM Sephadex™ C-25 column (GE Healthcare) equilibrated with MQ. The flow-through was discarded, and the column was subsequently washed with 12 column volumes (CV) of MQ. The peptide was eluted with 6 CV 2 M NaCl. The eluted peptide was then applied to a SIGMA-ALDRICH C18 Silica gel spherically equilibrated with 10 CV of 5% aq. MeCN containing 0.1% trifluoroacetic acid. After washing with a 10 CV of 5% aq. MeCN containing 0.1% trifluoroacetic acid, the peptide was eluted from the column using up to 10 CV of 50% aq. MeCN containing 0.1% trifluoroacetic acid. Fractions containing the eluted peptide were freeze-dried and dissolved in MQ water. After filtration through a 0.2 μm filter, the cationic peptides were purified on an Agilent 1260 Infinity HPLC system with a Phenomenex Aeris™ C18 column (250 × 4.6 mm, 3.6 μm particle size, 100 Å pore size). Acetonitrile was used as the mobile phase, and a gradient of 30-45% aq. MeCN over 30 min at 1 mL per min was used for separation. The purified lipo-tridecapeptides were eluted between 35 to 40 % aq. MeCN. The yields for brevibacillin, brevibacillin V, brevibacillin 2V and brevibacillin I were 5 mg/mL, 3 mg/mL, 1.5 mg/mL, 0.5 mg/mL, respectively.

### Mass Spectrometry

Matrix-assisted laser desorption ionization-time-of-flight (MALDI-TOF) mass spectrometer analysis was performed using a 4800 Plus MALDI TOF/TOF Analyzer (Applied Biosystems) in the linear-positive mode as in previous studies (Zhao et al., 2020a, 2020c, 2020b). Briefly, a 1 μL sample was spotted on the target and dried at room temperature. Subsequently, 0.6 μL of matrix solution (5 mg/mL of α-cyano-4-hydroxycinnamic acid) was spotted on each sample. After the samples had dried, MALDI-TOF MS was performed.

### LC-MS/MS Analysis

To gain insight into the molecular structures of the peptides, LC-MS/MS was performed, using a Q-Exactive mass spectrometer fitted with an Ultimate 3000 UPLC, an ACQUITY BEH C18 column (2.1 × 50 mm, 1.7 μm particle size, 200 Å; Waters), a HESI ion source, and a Orbitrap detector. A gradient of 5-90% MeCN with 0.1% formic acid (v/v) at a flow rate of 0.35 mL/min over 60 min was used. MS/MS was performed in a separate run in PRM mode, selecting the doubly and triply charged ion of the compound of interest.

### Minimum Inhibitory Concentration (MIC) Assay

MIC values were determined by using broth micro-dilution according to the standard guidelines (Wiegand et al., 2008). Briefly, the test medium was cation-adjusted Mueller-Hinton broth (MHB). Cell concentration was adjusted to approximately 5×10^5^ cells per ml. After 20 h of incubation at 37 °C, the MIC was defined as the lowest concentration of antibiotic with no visible growth (Zhao et al., 2020c). Each experiment was performed in triplicate.

### Hemolytic Activity

Hemolytic activity was determined with red blood cells freshly isolated from a healthy human. Peptides were added at final concentrations of 128, 64, 32, 16, 8, 4, 2, 1 and 0 μg /mL in PBS containing 2% (v/v) erythrocytes. The cells were incubated at 37 °C for 1 h and centrifuged for 5 min at 8,000 g. The supernatant was transferred to a 96-well plate, and the absorbance was measured at a wavelength of 450 nm with a Thermo Scientific Varioskan™ LUX multimode microplate reader. The absorbance relative to the positive control, which was treated with 10% Triton X-100, was defined as the percentage of hemolysis. The 50% Human blood cell hemolysis concentration (HC_50_) of peptides was calculated as previous studies described (Reed and Muench, 1938).

### Synergy Assays

The synergistic effect of brevibacillins with antibiotics was monitored using the checkerboard method (Doern, 2014). Briefly, antibiotics were two-fold series diluted while brevibacillins were added at a certain concentration (1, 2, or 4 μg/mL). Indicator strains were added at a final concentration of 5 × 10^5^ c.f.u/mL. After incubation at 37 ^0^C for 20 h, the OD_600_ of plates was determined. The fractional inhibitory concentration index (FICI) was calculated using the following formula FICI = (MICpeptide+antibiotic)/(MICpeptide) + (MICpeptide+antibiotic)/(MICantibiotic). The FICI value suggests a synergistic (≤ 0.5), addictive (> 0.5 - 1), no interaction (1 - 4), and antagonism (> 4) effect of the two compounds (Doern, 2014).

## Results

### Characterization of Peptides Discovered by Genome Mining

Many lipopeptide antibiotics have been discovered that originate from *Brevibacillus* spp (Barsby et al., 2001, 2006; Yang et al., 2016; Yang and Yousef, 2018; Wu et al., 2019). In a previous study, a cyclic lipopeptide, brevicidine, was discovered from *Brevibacillus laterosporus* DSM 25 by genome mining, which showed antimicrobial activity against Gram-negative pathogenic bacteria (Li et al., 2018; Zhao et al., 2020b). In the present study, we found by genome mining with the assistance of antiSMASH (Medema et al., 2011; Blin et al., 2019), that the genome of *Brevibacillus laterosporus* DSM 25 contains a gene cluster for synthesis of lipo-tridecapeptides (Figure 1 A and B). *Brevibacillus laterosporus* DSM 25 produced several cationic peptides that were purified by a CM SephadexTM C-25 column (GE Healthcare). After high-performance liquid chromatography (HPLC) purification (Supplementary Figure 1), MALDI-TOF MS was used to measure molecular weight of the purified peptides. Four compounds were found with a mass between 1550 to 1600 Da, i.e., compound **1** 1555 Da, compound **2** 1569 Da, compound **3** 1583 Da, and compound **4** 1597 Da (Supplementary Figures 1 and 2), which potentially belong to the lipo-tridecapeptide family. Further characterization of these peptides was performed by LC-MS/MS analysis, and the results are shown in Figure 2, Supplementary Figures 3, and 4. The structures of the four compounds were elucidated by using known lipo-tridecapeptides as reference templates (Barsby et al., 2001, 2006; Yang et al., 2016; Yang and Yousef, 2018; Wu et al., 2019). Two of these compounds are known peptides, i.e., brevibacillin (compound **3**) and brevibacillin V (compound **2**) (Yang et al., 2016; Wu et al., 2019), while the other two compounds are novel lipo-tridecapeptides that we named brevibacillin I (compound **4**) and brevibacillin 2V (compound **1**) (Figure 1C).

**Figure 1.**
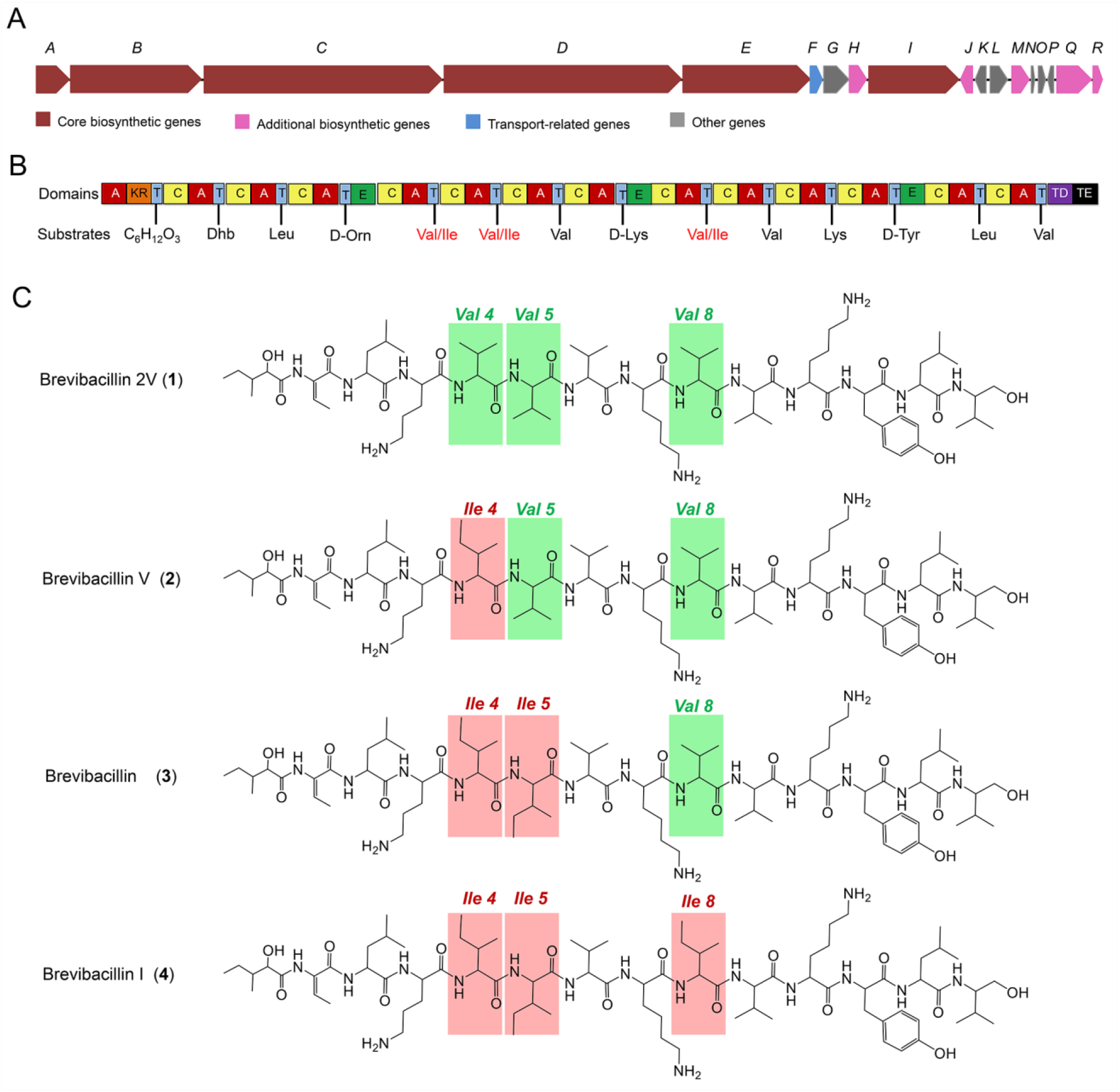
The structures of brevibacillins and the predicted biosynthetic gene cluster. **A**, The non-ribosomal peptide synthetases genes harbored by the *Brevibacillus laterosporus* DSM 25 genome; **B**, the catalytic domains encoded by the gene cluster, and the substrates incorporated by the respective modules. Domains: A, adenylation; KR, ketoreductase; T, thiolation; C, condensation; E, epimerization; TD, terminal reductase; TE, thioesterase; **C**, structures of brevibacillins. The red rectangles indicate the more hydrophobic amino acid residues of brevibacillin V, brevibacillin and brevibacillin I relative to brevibacillin 2V.

**Figure 2.**
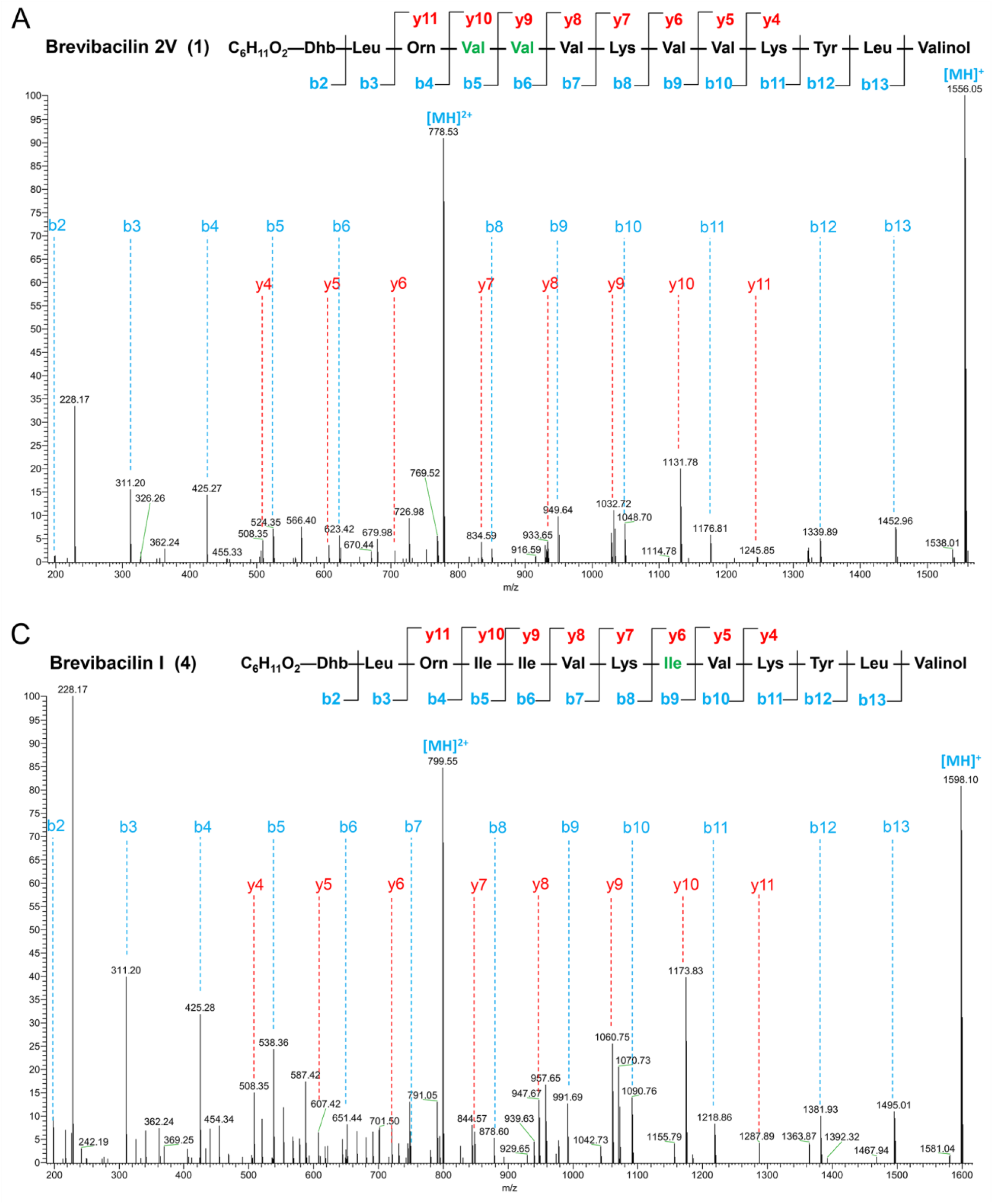
LC-MS/MS spectrums and the proposed structures of brevibacillin 2V (**A**) and brevibacillin I (**B**). Fragment ions are indicated.

### Brevibacillins Show Strong Antimicrobial Activity Against Gram-positive Bacterial Pathogens

The antimicrobial activity of brevibacillins against pathogenic bacteria was evaluated by a MIC assay. All of the brevibacillins showed strong antimicrobial activity against tested Gram-positive pathogenic bacteria, with a MIC value of 1-2 μg/mL, including difficult-to-treat antibiotic-resistant *Enterococcus faecium, Enterococcus faecalis* and *Staphylococcus aureus* (Table 1). The similar antimicrobial activities indicate that the varying amino acid residues in the different lipo-tridecapeptides have no significant influence on their antimicrobial activity. In addition, brevibacillins showed antimicrobial activity against Gram-negative pathogenic bacteria, yet these activities were much lower as compared to the antimicrobial activity against Gram-positive bacteria. These results are consistent with previous studies that show that lipo-tridecapeptides have antimicrobial activity against Gram-positive bacteria, but have insufficient antimicrobial activity against Gram-negative bacteria (Barsby et al., 2001, 2006; Yang et al., 2016; Wu et al., 2019). To assess the stability of brevibacillins towards proteolysis, a MIC assay was performed in the presence of 10% human blood plasma (Table 1). The results indicate that brevibacillins are very stable since the MIC values of them did not change by the addition of human blood plasma (Table 1).

**Table 1.** MIC values of brevibacillins against pathogenic bacteria.

| Microorganism | MIC (µg/mL) |  |  |  |  |  |
| --- | --- | --- | --- | --- | --- | --- |
|  | Bre | Bre V | Bre I | Bre 2V | Poly B | Nisin |
| **Gram-positive pathogenic bacteria** |  |  |  |  |  |  |
| *Staphylococcus aureus* ATCC15975 (MRSA) | 2 | 1-2 | 2 | 2 | 32 | 4-8 |
| *Enterococcus faecium* LMG16003 (VRE) | 2 | 2 | 2 | 2 | 16 | 4 |
| *Enterococcus faecium* LMG16003 (VRE)<br>plus 10% Human blood plasma | 2 | 2 | 2 | 2 | 16 | 4-8 |
| *Enterococcus faecalis* LMG16216 (VRE) | 2 | 2 | 2 | 2 | 16 | 4 |
| *Bacillus cereus* ATCC14579 | 1 | 1 | 2 | 2 | 8 | 4 |
| **Gram-negative pathogenic bacteria** |  |  |  |  |  |  |
| *Acinetobacter baumannii* ATCC17978 | 32 | 64 | 64 | 32 | 4 | 32 |
| *Escherichia coli* ATCC25922 | 32 | 32 | 32 | 16 | 2 | 64 |
| *Pseudomonas aeruginosa* LMG6395 | 64 | 64 | 64 | 64 | 1 | 128 |
| *Klebsiella pneumoniae* LMG20218 | 32 | 64 | 64 | 32 | 2 | 128 |

VRE, vancomycin resistant *Enterococcus*; MRSA, methicillin resistant *S. aureus*; Bre, brevibacillin ; Bre V, brevibacillin V; Bre I, brevibacillin I; Bre 2V, brevibacillin 2V; Poly B, polymyxin B.

### Brevibacillin 2V Does Not Exhibit Hemolytic Activity When Present at 128 µg/mL

To assess in an initial test the safety of brevibacillins to human beings or animals, the hemolytic activity of brevibacillins to human blood cells was monitored. Human blood cells were incubated in the presence of brevibacillins concentrations ranging from 1 μg/mL to 128 μg/mL. After incubation at 37 °C for 1 h, the OD_450_ of the supernatants was measured and the hemolytic activities of the brevibacillins were calculated as described in previous studies (Ling et al., 2015; Li et al., 2018). Brevibacillin, brevibacillin V and brevibacillin I showed significant hemolytic activity at concentrations that are 2-4 fold higher than their MIC values (Figure 3 and Table 1), with a half red blood cell hemolysis (HC_50_) concentration of 18.8 ± 0.5 μg/mL, 73.1 ± 1.4 μg/mL, and 25.1 ± 3.9 μg/mL, respectively. In addition, in the presence of 64 μg/mL of either brevibacillin/brevibacillin I or 128 μg/mL of brevibacillin V, human blood cells were completely lysed (Figure 3). In contrast, brevibacillin 2V showed no hemolytic activity against human red blood cells at the high concentration of 128 μg/mL (Figure 3). These results demonstrate that brevibacillin 2V shows high potential, justifying its further study and development as an alternative antibiotic for controlling antibiotic-resistant bacterial pathogens.

**Figure 3.**
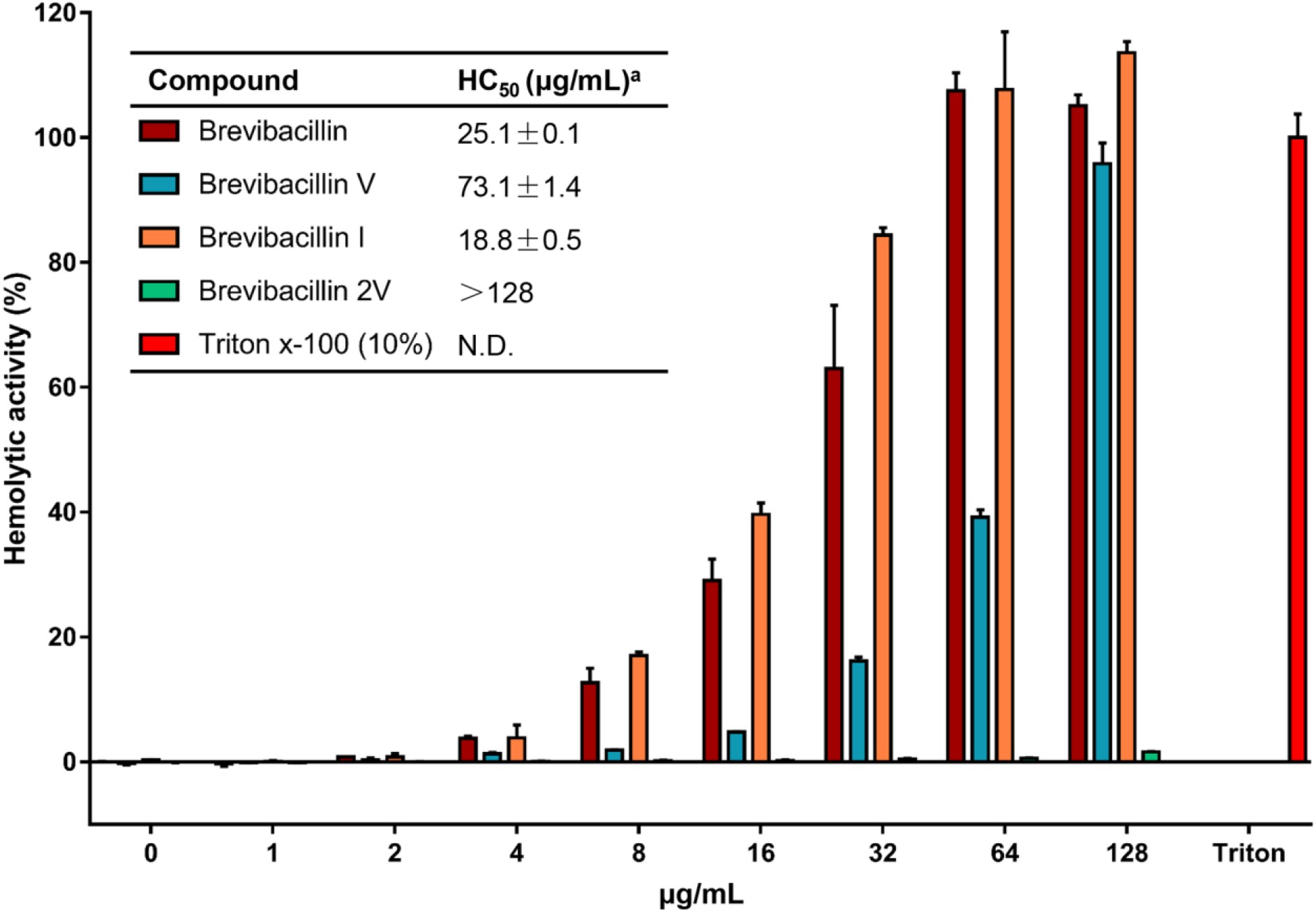
Brevibacillins activity against eukaryotic cells. Human erythrocytes were incubated with brevibacillins at concentrations ranging from 1 to 128 μg/mL. Their hemolytic activity was assessed by the release of hemoglobin. Cells treated without a tested compound were used as no lysis control. Cells treated with 10% Triton X-100 were used as complete lysis control. The data are representative of three independent experiments. ^a^ HC_50_ is the concentration that causes 50% hemolysis of Human red blood cells. N.D., not determined.

### Brevibacillins Show Synergistic Activity with Other Antibiotics Against Gram-negative Pathogenic Bacteria

Brevibacillins show strong antimicrobial activity against Gram-positive bacterial pathogens, but have insufficient antimicrobial activity against Gram-negative bacterial pathogens (Table 1). To expand the antimicrobial potential of brevibacillins, we investigated the synergistic activity of brevibacillins with some antibiotics (Nalidixic acid, Azithromycin, Rifampicin, and Amikacin) (Moore, 2015) against Gram-negative pathogens. The synergistic effect of peptides was evaluated by determining a fractional inhibitory concentration index (FICI) value, which can indicate either a synergistic (≤ 0.5), addictive (> 0.5 - 1), no interaction (1 - 4), or antagonistic (> 4) effect of the two compounds (Doern, 2014). The results are shown in Table 2 and Supplementary Tables 2, 3, and 4. Brevibacillin, brevibacillin V, brevibacillin I and brevibacillin 2V showed synergistic activity with the tested antibiotics against corresponding Gram-negative pathogens. Most importantly, brevibacillins showed the highest synergistic activity with Amikacin against *A. baumannii* in all tests (Table 2 and Supplementary Tables 2, 3, and 4), with *A. baumannii* being one of the three critical priority pathogens for R&D of new antibiotics (Organization, 2017). The MIC value of Amikacin against *A. baumannii* decreased 32-64 folds in the presence of 4μg/mL brevibacillins (Table 2 and Supplementary Tables 2, 3, and 4). These results demonstrate that brevibacillins (at least brevibacillin 2V) could be used as adjuvant for other antibiotics to treat Gram-negative bacterial pathogen infections.

**Table 2.**
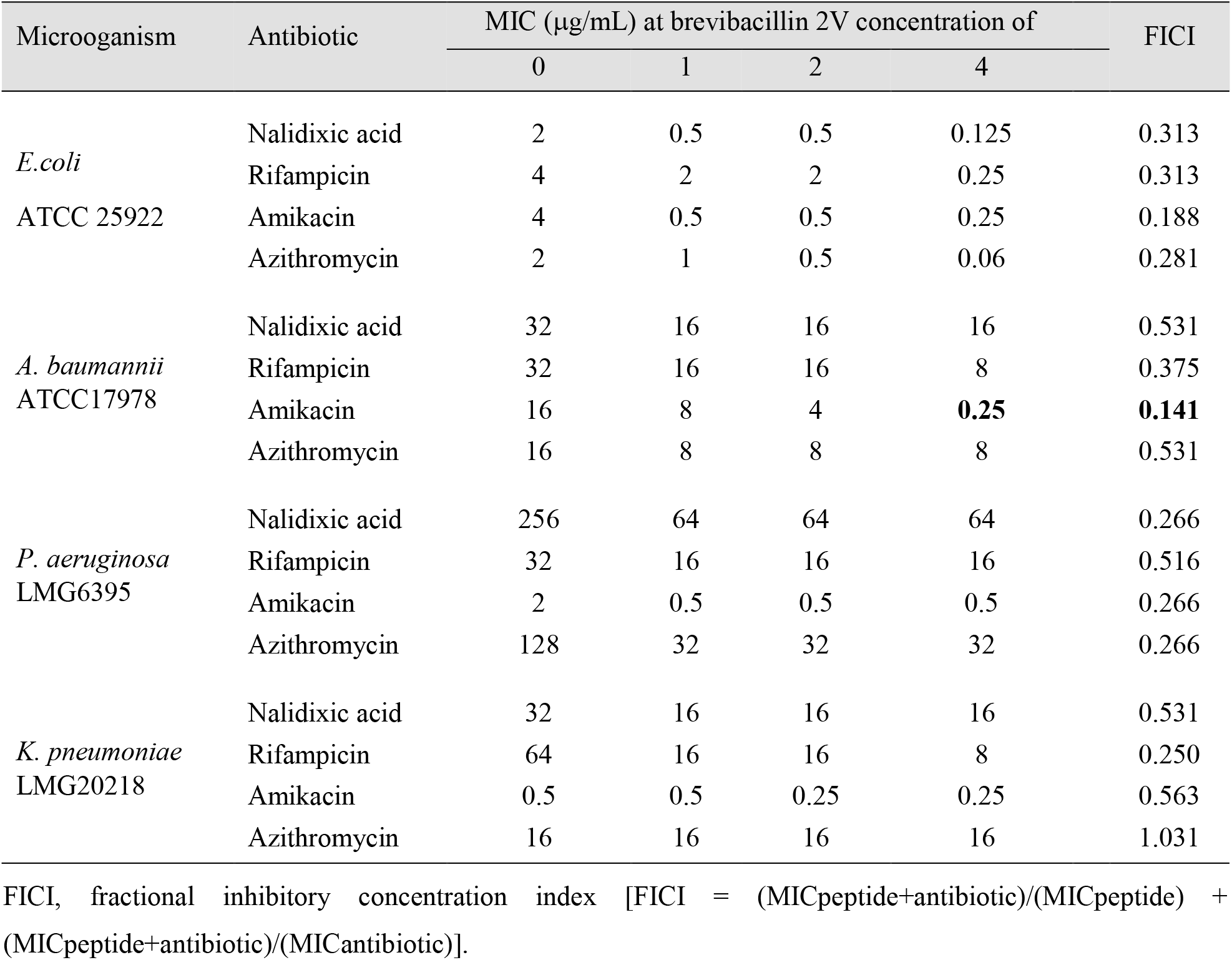
Synergistic effect between brevibacillin 2V and antibiotics.

## Discussion

Genome mining is a practical approach for discovering bioactive natural products from microorganisms (Montalbán-López et al., 2021). Genome mining tools with different specific uses have been developed, including BAGEL4 (van Heel et al., 2018), antiSMASH (Blin et al., 2019), and RiPP-PRISM (Skinnider et al., 2015). AntiSMASH is a well-known genome mining tool for discovering NRPs with antimicrobial activity (Blin et al., 2019). For instance, two non-ribosomally produced cyclic lipopeptides, i.e., brevicidine and laterocidine, were discovered by genome mining with the assistance of antiSMASH (Li et al., 2018). In this study, a lipo-tridecapeptides synthetic gene cluster was discovered from *Brevibacillus laterosporus* DSM 25 by genome mining with the assistance of antiSMASH (Medema et al., 2011; Blin et al., 2019). Subsequently, antimicrobial lipo-tridecapeptides were isolated and identified by following the pipeline in Figure 4.

**Figure 4.**
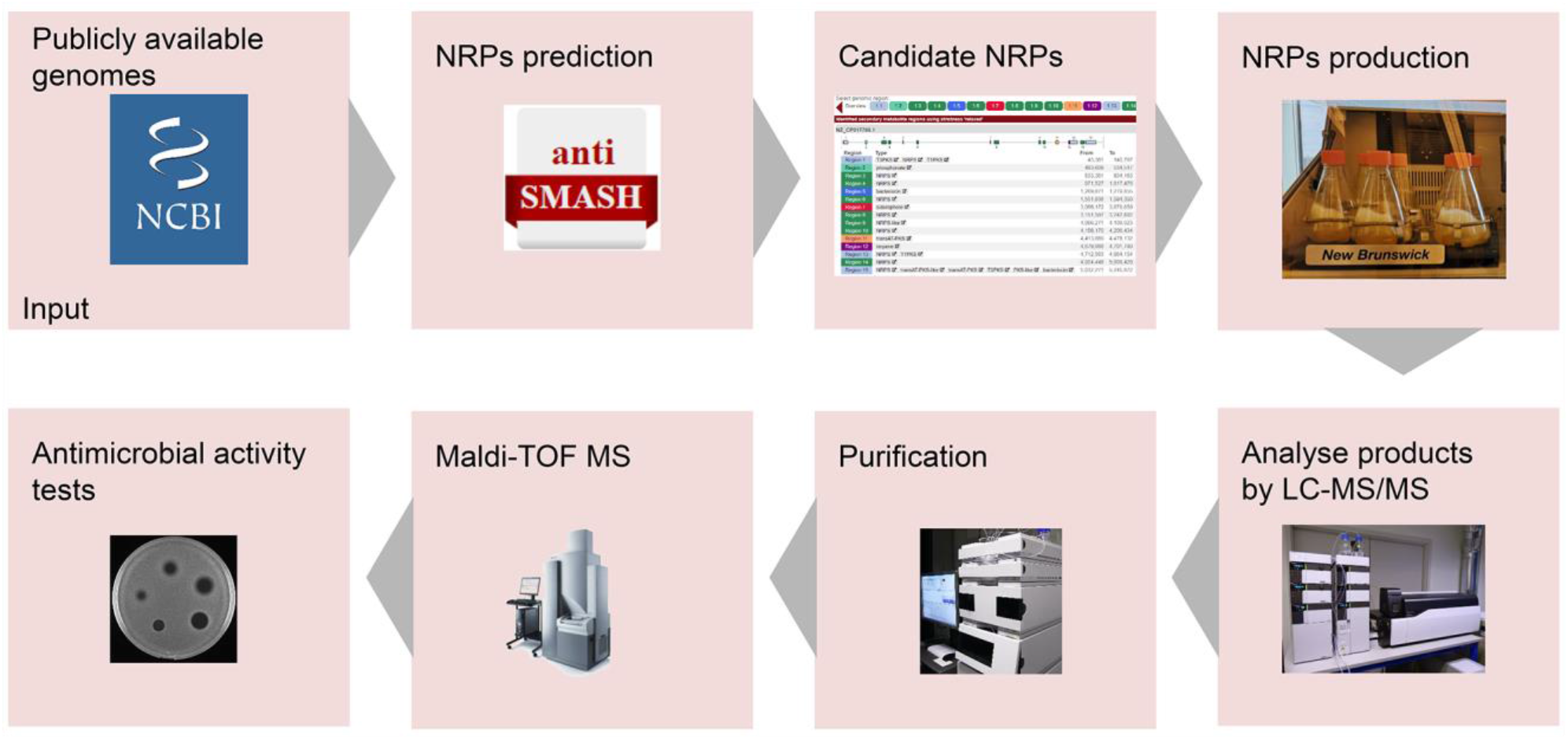
Pipeline for the development of novel NRP-based antimicrobials. Following the arrows, the process starts with the identification of NRP clusters. Next, the predicted NRPs have to be produced, isolated, and characterized. Finally, the antimicrobial activity of purified NRPs could be tested.

In this study, we isolated the predicted lipo-tridecapeptides from the supernatant of *Brevibacillus laterosporus* DSM 25 culture. This allows an iron change-based CM SephadexTM C-25 column to be applied to purify the produced products since lipo-tridecapeptides are cationic peptides. In contrast, lipo-tridecapeptides were previously isolated from bacterial cells (Barsby et al., 2001, 2006; Yang et al., 2016), which is more difficult to process. Thus, the here found lipo-tridecapeptides are much more readily purified than previously reported lipo-tridecapeptides. For elucidation of the purified lipo-tridecapeptides structures, we used previously reported lipo-tridecapeptides as templates. Such template-based structure elucidation approach has been successfully used in previous studies to elucidate peptide structures (Barsby et al., 2006; Wu et al., 2019). These results suggest that a template-based structure elucidation is a helpful approach for elucidating the structures of peptides that belong to the same type. All structures were confirmed by MS.

Many lipo-tridecapeptides have shown antimicrobial activity against pathogenic bacteria, including antibiotic-resistant *Staphylococcus aureus* and *Enterococcus spp*. (Barsby et al., 2006; Yang et al., 2016; Wu et al., 2019). However, the high hemolytic activity of lipo-tridecapeptides has limited their potential for being developed as therapeutics (Figure 3) (Li et al., 2018). In this study, one of the discovered novel lipo-tridecapeptides, brevibacillin 2V, showed no hemolytic activity at the high concentration of 128 μg/mL (Figure 3). Moreover, the hemolytic activity of brevibacillins shows a positive correlation with their calculated hydrophobicity, i.e., brevibacillin I > brevibacillin > brevibacillin V > brevibacillin 2V (Figure 1C and Supplementary Figure S1). These results demonstrate that brevibacillin 2V shows a high potential to be further studied and developed as an alternative antibiotic for controlling antibiotic-resistant bacterial pathogens. In addition, the positive correlation of hemolytic activity and hydrophobicity provides a guideline for the synthesis of lipo-tridecapeptide analogs in future studies. The use of peptides as adjuvants for marketed antibiotics has attracted the attention of several researchers. Cochrane et al. reported that unacylated tridecaptin A1 acts as an effective sensitizer of Gram-negative bacteria to other antibiotics (Cochrane and Vederas, 2014). More recently, Li et al. reported that outer-membrane-acting peptides showed synergistic activity with lipid II-targeting antibiotics against Gram-negative pathogens (Li et al., 2021). In the present study, brevibacillins showed synergistic activity with marketed antibiotics against Gram-negative pathogens. These results suggest that more antibiotics can be tested for developing brevibacillins (at least brevibacillin 2V) as adjuvants for other antibiotics to treat Gram-negative bacterial pathogen infections.

In conclusion, in this study, we characterized a novel lipo-tridecapeptide, i.e., brevibacillin 2V, which was identified from *Brevibacillus laterosporus* DSM 25 by genome mining and subsequently isolated, and purified. Brevibacillin 2V has strong antimicrobial activity against antibiotic-resistant Gram-positive bacterial pathogens, and it shows effective synergistic activity with other antibiotics against Gram-negative bacterial pathogens. Notably, brevibacillin 2V has a much lower hemolytic activity towards eukaryotic cells than previously reported NRPs of the lipo-tridecapeptide family, including all known brevibacillins, which makes it a promising candidate for antibiotic development. This study provides a novel and promising antibiotic candidate (brevibacillin 2V) with low hemolytic activity, which can be used either directly as it is, or serves as a template for further total synthesis and modifications.

## Data Availability Statement

All datasets presented in this study are included in the the article/**Supplementary Material**.

## Author Contributions

Oscar P Kuipers and Xinghong Zhao conceived the project. Xinghong Zhao designed and carried out the experiments, analyzed data, and wrote the manuscript. Oscar P Kuipers and Eefjan Breukink supervised the work and corrected the manuscript. Xiaoqi Wang, Rhythm Shukla, Raj Kumar, and Markus Weingarth did experimental work. All authors contributed to and commented on the manuscript text and approved its final version.

## Acknowledgments

We thank Marcel P. de Vries (University of Groningen) for support with LC-MS/MS.

## References

Barsby, T., Kelly, M. T., Gagné, S. M., and Andersen, R. J. (2001). Bogorol A produced in culture by a marine Bacillus sp. reveals a novel template for cationic peptide antibiotics. Org. Lett. 3, 437–440.

Barsby, T., Warabi, K., Sørensen, D., Zimmerman, W. T., Kelly, M. T., and Andersen, R. J. (2006). The bogorol family of antibiotics: template-based structure elucidation and a new approach to positioning enantiomeric pairs of amino acids. J. Org. Chem. 71, 6031–6037.

Blin, K., Shaw, S., Steinke, K., Villebro, R., Ziemert, N., Lee, S. Y., et al. (2019). antiSMASH 5.0: updates to the secondary metabolite genome mining pipeline. Nucleic Acids Res. 47, W81–W87.

Cochrane, S. A., Lohans, C. T., Brandelli, J. R., Mulvey, G., Armstrong, G. D., and Vederas, J. C. (2014). Synthesis and structure–activity relationship studies of N-terminal analogues of the antimicrobial peptide tridecaptin A1. J. Med. Chem. 57, 1127–1131.

Cochrane, S. A., and Vederas, J. C. (2014). Unacylated tridecaptin A1 acts as an effective sensitiser of Gram-negative bacteria to other antibiotics. Int. J. Antimicrob. Agents 44, 493–499.

Cociancich, S., Pesic, A., Petras, D., Uhlmann, S., Kretz, J., Schubert, V., et al. (2015). The gyrase inhibitor albicidin consists of p-aminobenzoic acids and cyanoalanine. Nat. Chem. Biol. 11, 195.

Doern, C. D. (2014). When does 2 plus 2 equal 5? A review of antimicrobial synergy testing. J. Clin. Microbiol. 52, 4124–4128.

Hamamoto, H., Urai, M., Ishii, K., Yasukawa, J., Paudel, A., Murai, M., et al. (2015). Lysocin E is a new antibiotic that targets menaquinone in the bacterial membrane. Nat. Chem. Biol. 11, 127.

Li, Q., Cebrián, R., Montalbán-López, M., Ren, H., Wu, W., and Kuipers, O. P. (2021). Outer-membrane-acting peptides and lipid II-targeting antibiotics cooperatively kill Gram-negative pathogens. Commun. Biol. 4, 1–11.

Li, Y.-X., Zhong, Z., Zhang, W.-P., and Qian, P.-Y. (2018). Discovery of cationic nonribosomal peptides as Gram-negative antibiotics through global genome mining. Nat. Commun. 9, 3273.

Ling, L. L., Schneider, T., Peoples, A. J., Spoering, A. L., Engels, I., Conlon, B. P., et al. (2015). A new antibiotic kills pathogens without detectable resistance. Nature 517, 455.

Medema, M. H., Blin, K., Cimermancic, P., de Jager, V., Zakrzewski, P., Fischbach, M. A., et al. (2011). antiSMASH: rapid identification, annotation and analysis of secondary metabolite biosynthesis gene clusters in bacterial and fungal genome sequences. Nucleic Acids Res. 39, W339–W346.

Montalbán-López, M., Scott, T. A., Ramesh, S., Rahman, I. R., van Heel, A. J., Viel, J. H., et al. (2021). New developments in RiPP discovery, enzymology and engineering. Nat. Prod. Rep. 38, 130–239. doi:10.1039/d0np00027b.

Moore, D. (2015). Antibiotic classification and mechanism. Retrieved August 24.

Organization, W. H. (2017). Prioritization of pathogens to guide discovery, research and development of new antibiotics for drug-resistant bacterial infections, including tuberculosis. World Health Organization.

Reed, L. J., and Muench, H. (1938). A SIMPLE METHOD OF ESTIMATING FIFTY PER CENT ENDPOINTS,. Am. J. Epidemiol. 27.

Skinnider, M. A., Dejong, C. A., Rees, P. N., Johnston, C. W., Li, H., Webster, A. L. H., et al. (2015). Genomes to natural products prediction informatics for secondary metabolomes (PRISM). Nucleic Acids Res. 43, 9645–9662.

Süssmuth, R. D., and Mainz, A. (2017). Nonribosomal peptide synthesis—principles and prospects. Angew. Chemie Int. Ed. 56, 3770–3821.

van Heel, A. J., de Jong, A., Song, C., Viel, J. H., Kok, J., and Kuipers, O. P. (2018). BAGEL4: a user-friendly web server to thoroughly mine RiPPs and bacteriocins. Nucleic Acids Res. 46, W278–W281.

Wiegand, I., Hilpert, K., and Hancock, R. E. W. (2008). Agar and broth dilution methods to determine the minimal inhibitory concentration (MIC) of antimicrobial substances. Nat. Protoc. 3, 163.

Wu, Y., Zhou, L., Lu, F., Bie, X., Zhao, H., Zhang, C., et al. (2019). Discovery of a Novel Antimicrobial Lipopeptide, Brevibacillin V, from Brevibacillus laterosporus fmb70 and Its Application on the Preservation of Skim Milk. J. Agric. Food Chem. 67, 12452–12460.

Yang, X., Huang, E., Yuan, C., Zhang, L., and Yousef, A. E. (2016). Isolation and structural elucidation of brevibacillin, an antimicrobial lipopeptide from Brevibacillus laterosporus that combats drug-resistant Gram-positive bacteria. Appl. Environ. Microbiol. 82, 2763–2772.

Yang, X., and Yousef, A. E. (2018). Antimicrobial peptides produced by Brevibacillus spp.: structure, classification and bioactivity: a mini review. World J. Microbiol. Biotechnol. 34, 57.

Zhao, X., Cebrián, R., Fu, Y., Rink, R., Bosma, T., Moll, G. N., et al. (2020a). High-Throughput Screening for Substrate Specificity-Adapted Mutants of the Nisin Dehydratase NisB. ACS Synth. Biol. 9, 1468–1478. doi:10.1021/acssynbio.0c00130.

Zhao, X., Li, Z., and Kuipers, O. P. (2020b). Mimicry of a Non-ribosomally Produced Antimicrobial, Brevicidine, by Ribosomal Synthesis and Post-translational Modification. Cell Chem. Biol. 27, 1262–1271.e4. doi:10.1016/j.chembiol.2020.07.005.

Zhao, X., Yin, Z., Breukink, E., Moll, G. N., and Kuipers, O. P. (2020c). An Engineered Double Lipid II Binding Motifs-Containing Activity against Enterococcus faecium. Antimicrob. Agents Chemother. 64, 1–12.

