## Supplemental Figures and Tables for "Brevibacillin 2V, a novel antimicrobial lipopeptide with an exceptional low hemolytic activity"

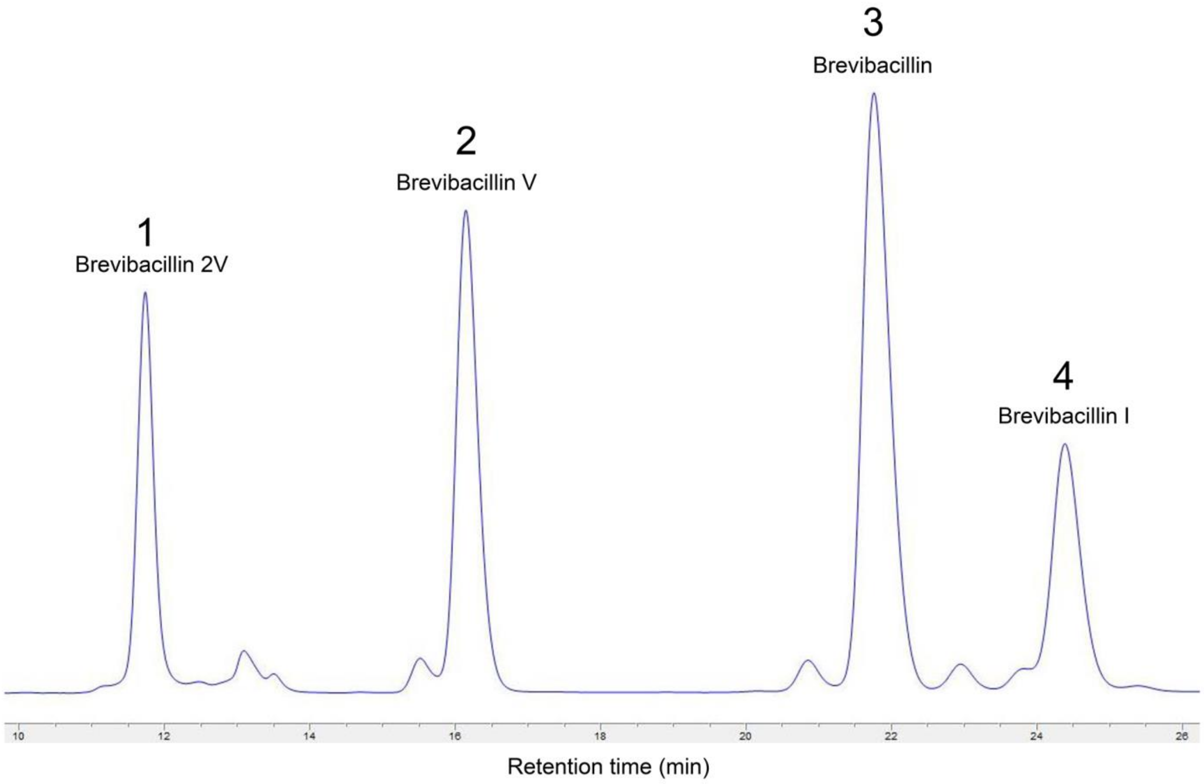


**Supplementary Figure 1.** The HPLC spectrum of brevibacillins. The longer retention time correlated with the higher hydrophobicity of brevibacillins, which shows hydrophobicity that brevibacillin I ＞ brevibacillin＞ brevibacillin V ＞ brevibacillin 2V. Compounds 1, 2, 3 and 4 were elucidated by further studies as brevibacillin 2V, brevibacillin V, brevibacillin and brevibacillin I, respectively.

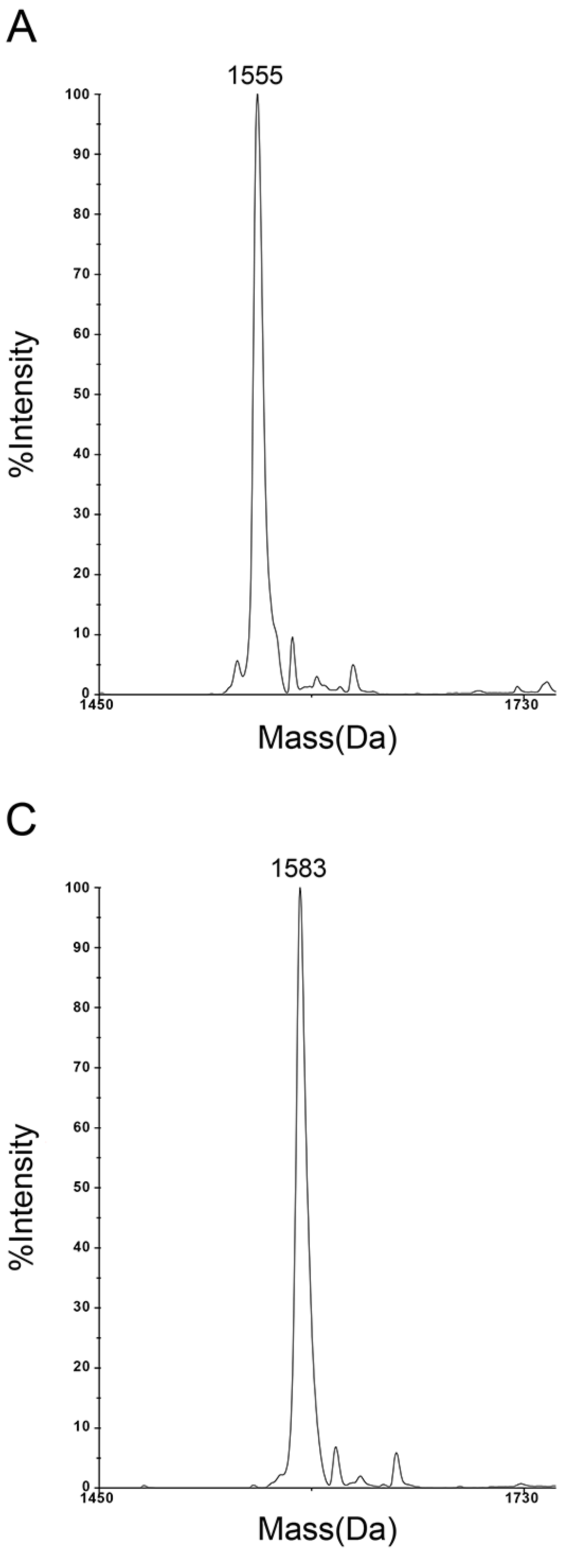


**Supplementary Figure 2.** MALDI-TOF spectrum of HPLC purified brevibacillins. **A**, brevibacillin 2V; **B**, brevibacillin V; **C**, brevibacillin; **D**, brevibacillin I.

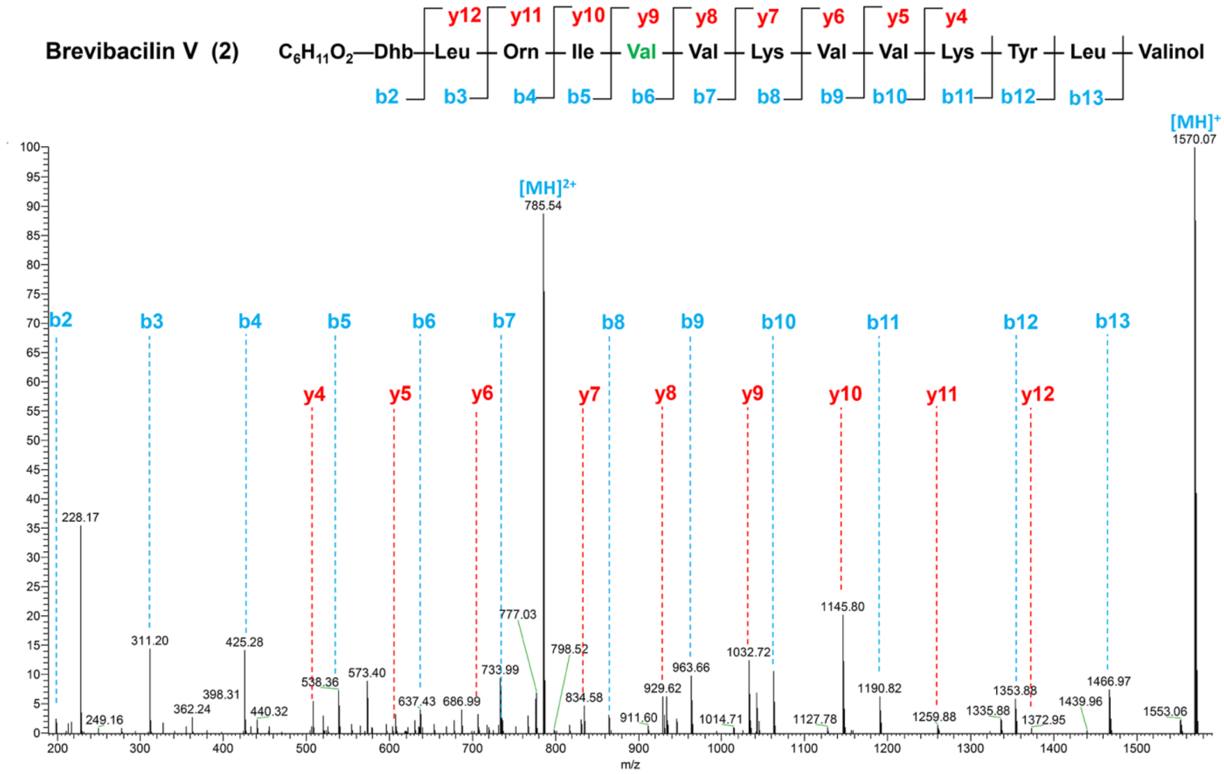


**Supplementary Figure 3.** LC-MS/MS spectrum and the proposed structure of brevibacillin V (2). Fragment ions are indicated.

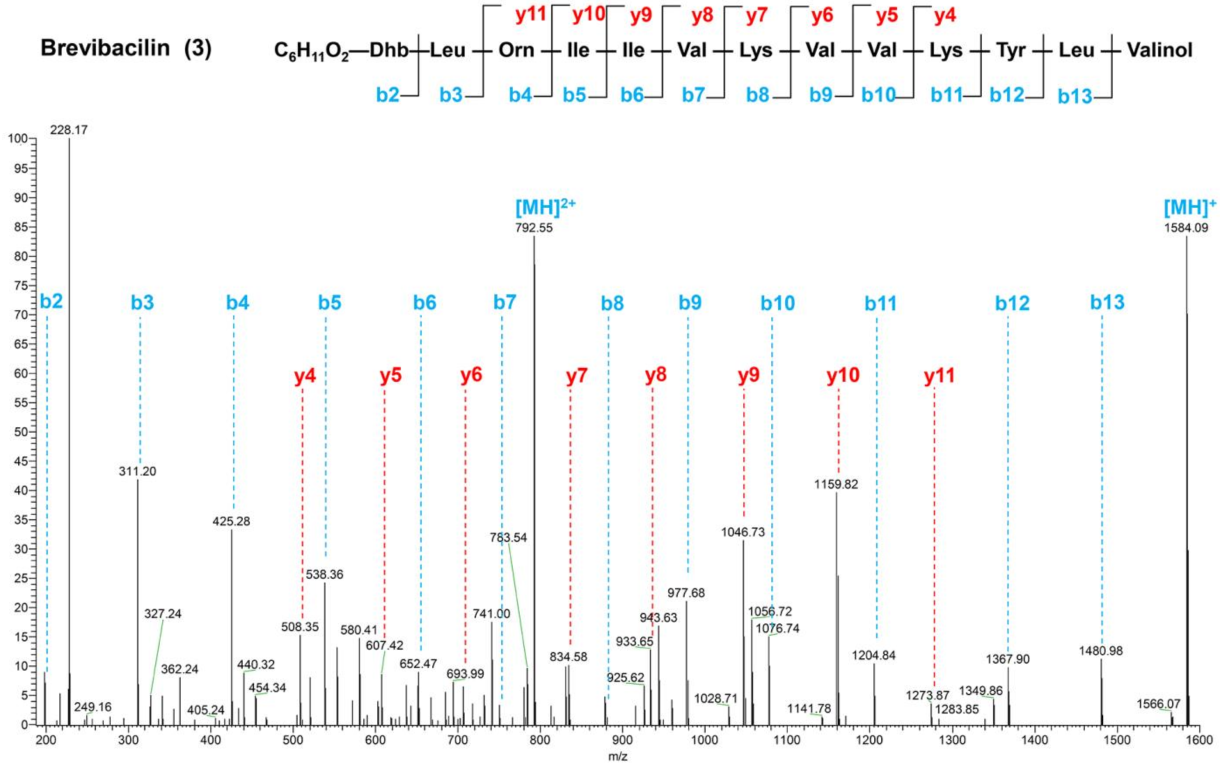


**Supplementary Figure 4.** LC-MS/MS spectrum and the proposed structure of brevibacillin (3). Fragment ions are indicated.

**Supplementary Table 1** Stains used in this study.

| Strains | Characteristics and purpose |
|---|---|
| *Brevibacillus laterosporus* | DSM 25, host strain for production of brevibacillin |
| *Bacillus cereus* | ATCC14579, indicator strain |
| *Enterococcus faecalis* | LMG16216 (VRE), indicator strain |
| *Staphylococcus aureus* | ATCC15975 (MRSA), indicator strain |
| *Enterococcus faecium* | LMG16003 (VRE), indicator strain |
| *Acinetobacter baumannii* | ATCC17978, indicator strain |
| *Escherichia coli* | ATCC25922, indicator strain |
| *Klebsiella pneumoniae* | LMG20218, indicator strain |
| *Pseudomonas aeruginosa* | LMG6395, indicator strain |

**Supplementary Table 2** Synergistic effect between brevibacillin V and antibiotics.

| Microoganism | Antibiotic | MIC(μg/mL) at brevibacillin V concentrations of | | | | FICI |
|---|---|---|---|---|---|---|
| | | 0 | 1 | 2 | 4 | |
| | Nalidixic acid | 2 | 2 | 1 | 0.5 | 0.375 |
| *E.coli* | Rifampicin | 4 | 4 | 2 | 1 | 0.375 |
| ATCC 25922 | Amikacin | 4 | 1 | 1 | 0.5 | 0.250 |
| | Azithromycin | 2 | 2 | 1 | 1 | 0.563 |
| | Nalidixic acid | 32 | 32 | 32 | 32 | 1.016 |
| *A. baumannii* | Rifampicin | 32 | 16 | 16 | 16 | 0.516 |
| ATCC17978 | Amikacin | 16 | 4 | 2 | **0.5** | **0.094** |
| | Azithromycin | 16 | 16 | 16 | 16 | 1.016 |
| | Nalidixic acid | 256 | 128 | 128 | 128 | 0.516 |
| *P. aeruginosa* | Rifampicin | 32 | 16 | 16 | 16 | 0.516 |
| LMG6395 | Amikacin | 2 | 0.5 | 0.5 | 0.5 | 0.266 |
| | Azithromycin | 128 | 64 | 64 | 64 | 0.516 |
| | Nalidixic acid | 32 | 32 | 32 | 32 | 1.016 |
| *K. pneumoniae* | Rifampicin | 64 | 16 | 16 | 16 | 0.266 |
| LMG20218 | Amikacin | 0.5 | 0.5 | 0.5 | 0.5 | 1.016 |
| | Azithromycin | 16 | 16 | 16 | 16 | 1.016 |

FICI, fractional inhibitory concentration index [FICI = (MICpeptide+antibiotic)/(MICpeptide) + (MICpeptide+antibiotic)/(MICantibiotic)].

**Supplementary Table 3** Synergistic effect between brevibacillin and antibiotics.

| Microoganism | Antibiotic | MIC (μg/mL) at brevibacillin concentrations of | | | | FICI |
|---|---|---|---|---|---|---|
| | | 0 | 1 | 2 | 4 | |
| | Nalidixic acid | 2 | 1 | 1 | 0.5 | 0.375 |
| *E.coli* | Rifampicin | 4 | 2 | 1 | 0.5 | 0.250 |
| ATCC 25922 | Amikacin | 4 | 0.5 | 0.5 | 0.5 | 0.156 |
| | Azithromycin | 2 | 1 | 0.5 | 0.125 | 0.188 |
| | Nalidixic acid | 32 | 32 | 32 | 16 | 0.625 |
| *A. baumannii* | Rifampicin | 32 | 8 | 8 | 8 | 0.281 |
| ATCC17978 | Amikacin | 16 | 2 | 0.5 | **0.25** | **0.094** |
| | Azithromycin | 16 | 8 | 8 | 8 | 0.531 |
| | Nalidixic acid | 256 | 128 | 128 | 128 | 0.516 |
| *P. aeruginosa* | Rifampicin | 32 | 16 | 8 | 8 | 0.281 |
| LMG6395 | Amikacin | 2 | 0.5 | 0.5 | 0.5 | 0.266 |
| | Azithromycin | 128 | 64 | 64 | 64 | 0.516 |
| | Nalidixic acid | 32 | 32 | 16 | 16 | 0.563 |
| *K. pneumoniae* | Rifampicin | 64 | 16 | 16 | 16 | 0.281 |
| LMG20218 | Amikacin | 0.5 | 0.5 | 0.5 | 0.5 | 1.031 |
| | Azithromycin | 16 | 16 | 16 | 16 | 1.031 |

FICI, fractional inhibitory concentration index [FICI = (MICpeptide+antibiotic)/(MICpeptide) + (MICpeptide+antibiotic)/(MICantibiotic)].

**Supplementary Table 4** Synergistic effect between brevibacillin I and antibiotics.

| Microoganism | Antibiotic | MIC (μg/mL) at brevibacillin I concentrations of | | | | FICI |
|---|---|---|---|---|---|---|
| | | 0 | 1 | 2 | 4 | |
| *E.coli* ATCC 25922 | Nalidixic acid | 2 | 1 | 0.5 | 0.5 | 0.313 |
| | Rifampicin | 4 | 2 | 2 | 2 | 0.531 |
| | Amikacin | 4 | 1 | 1 | 1 | 0.281 |
| | Azithromycin | 2 | 1 | 1 | 0.5 | 0.375 |
| *A. baumannii* ATCC17978 | Nalidixic acid | 32 | 16 | 16 | 16 | 0.516 |
| | Rifampicin | 32 | 16 | 16 | 16 | 0.516 |
| | Amikacin | 16 | 8 | 4 | **0.25** | **0.078** |
| | Azithromycin | 16 | 8 | 8 | 8 | 0.516 |
| *P. aeruginosa* LMG6395 | Nalidixic acid | 256 | 64 | 64 | 64 | 0.266 |
| | Rifampicin | 32 | 16 | 16 | 16 | 0.516 |
| | Amikacin | 2 | 0.5 | 0.5 | 0.5 | 0.266 |
| | Azithromycin | 128 | 32 | 32 | 32 | 0.266 |
| *K. pneumoniae* LMG20218 | Nalidixic acid | 32 | 16 | 16 | 16 | 0.516 |
| | Rifampicin | 64 | 16 | 16 | 16 | 0.266 |
| | Amikacin | 0.5 | 0.5 | 0.5 | 0.25 | 0.563 |
| | Azithromycin | 16 | 16 | 16 | 16 | 1.016 |

FICI, fractional inhibitory concentration index [FICI = (MICpeptide+antibiotic)/(MICpeptide) + (MICpeptide+antibiotic)/(MICantibiotic)].
